# EcoFun-MAP: An Ecological Function Oriented Metagenomic Analysis Pipeline

**DOI:** 10.1101/2022.04.05.481366

**Authors:** Zhou Jason Shi, Naijia Xiao, Daliang Ning, Renmao Tian, Ping Zhang, Daniel Curtis, Joy D. Van Nostrand, Liyou Wu, Terry C. Hazen, Andrea M. Rocha, Zhili He, Adam P. Arkin, Mary K. Firestone, Jizhong Zhou

## Abstract

Annotating ecological functions of environmental metagenomes is challenging due to a lack of specialized reference databases and computational barriers. Here we present the **Eco**logical **Fun**ction oriented **M**etagenomic **A**nalysis **P**ipeline (EcoFun-MAP) for efficient analysis of shotgun metagenomes in the context of ecological functions. We manually curated a reference database of EcoFun-MAP which is used for GeoChip design. This database included ∼1,500 functional gene families that were catalogued by important ecological functions, such as carbon, nitrogen, phosphorus, and sulfur cycling, metal homeostasis, stress responses, organic contaminant degradation, antibiotic resistance, microbial defense, electron transfer, virulence and plant growth promotion. EcoFun-MAP has five optional workflows from ultra-fast to ultra-conservative, fitting different research needs from functional gene exploration to stringent comparison. The pipeline is deployed on High Performance Computing (HPC) infrastructure with a highly accessible web-based interface. We showed that EcoFun-MAP is accurate and can process multi-million short reads in a minute. We applied EcoFun-MAP to analyze metagenomes from groundwater samples and revealed interesting insights of microbial functional traits in response to contaminations. EcoFun-MAP is available as a public web server at http://iegst1.rccc.ou.edu:8080/ecofunmap/.

## Introduction

High throughput sequencing (HTS) and associated genomic technologies have revolutionized microbial ecology studies^1–5^ in the past decade. It allows in-depth profiling of microbial communities from environmental samples and leads to novel insights into microbial species, metabolic functions and pathways^6–8^. Shotgun metagenomic sequencing is one major application of HTS technology for microbial ecology studies^9^. It randomly recovers short or long reads from metagenomes, avoids selective amplification and thus several related biases and limitations in amplicon sequencing^10–12^, and has the potential to accurately quantify the abundances of both microbial taxa and functional genes^13^.

Shotgun metagenomic sequencing often generates a large volume of data and requires intensive complicated computational analysis to survey a typical environmental metagenome, e.g. soil metagenome. To meet the data analysis needs, many computational resources have been made available. Those resources have three main categories, including standalone programs for individual steps (e.g. quality control^14–16^, assembly^17,18^, gene prediction^19,20^, and sequence alignment^21–23^, reference databases^24–28^ and integrated analysis pipelines^29–32^).

Regardless of the available resources, we still face challenges for annotating functional genes of metagenomes that are ecologically important. First, most reference databases, e.g., NCBI nr, are for general purpose and lack focus on ecological functions. Using those databases without further filtration/distillation could result in unnecessary computing and data interpretation difficulties. Second, lack of computing skills and advanced hardware resources is still prevalent among microbial ecologists, which hinders the use of standalone programs and databases, especially those with complex interfaces and insufficient documentation. Integrated analysis pipelines, particularly web-based applications, which provide universal access and require little computing skills, are uniquely positioned to address this challenge. Nevertheless, few pipelines of such are available, efficient or ecology-oriented.

Here we developed an **Eco**logical **Fun**ction oriented **M**etagenomics **A**nalysis **P**ipeline (EcoFun-MAP), which uses a gene-centric paradigm to ease functional analysis of metagenomes. We manually curated the reference database of EcoFun-MAP by selecting and categorizing a comprehensive collection of microbial genes that were important to ecological functions and geochemical processes, which are used to design comprehensive GeoChip – a high throughput array-based technology for dissecting microbial community functions important to biogeochemistry, ecology, environmental sciences, agriculture, as well as human health^33–36^. The database included both relevant nucleotide and amino acid sequences, and hidden Markov models that are necessary for computing facilitation. EcoFun-MAP is also designed with several distinct data analysis workflows and evaluated for both speed and accuracy. To promote efficiency and accessibility, EcoFun-MAP was deployed on High-Performance Computing (HPC) infrastructure with a web-based user interface. In addition, we applied EcoFun-MAP to analyze metagenomes from groundwater samples and demonstrated its high effectiveness in revealing compositional variations of relevant functional genes along the contamination gradient.

## Results

### Overview of EcoFun-MAP

EcoFun-MAP is a fully automated pipeline that performs efficient functional gene annotation of sequencing reads from shotgun metagenomes. It consists of a reference database of functional genes and several computing workflows for gene annotation.

The reference database of EcoFun-MAP has four main modules, including a DIAMOND^23^ index of seed sequences (EFM-DI-DB-S), gene family Hidden Markov Models (EFM-HMM-DB), a DIAMOND (EFM-DI-DB-R) and NCBI-BLAST^37^ (EFM-BLAST-DB) index of full sequences (Fig. 1). To build these modules, we first manually selected the protein seed sequences for all functional gene families, which are pooled to make the EFM-DI-DB-S. We aligned seed sequences of each functional gene family and used the resulting multiple sequence alignment to build the EFM-HMM-DB. Both EFM-DI-DB-R and EFM-BLAST-DB modules use full reference sequences rather than the seed sequences only. We downloaded a large number of candidate sequences with keyword-based queries from NCBI GenBank^25^. Although our queries were crafted carefully, there might still be some false sequences in the candidate sequences due to possible mis-annotations. To exclude these false sequences, we used an iterative HMM searching procedure, which has the following steps: (i) set up an initial e-value cutoff = 10, (ii) searching the candidate sequences against the EFM-HMM-DB using the e-value cutoff, (iii) manually evaluate the resulting candidate sequences passed the cutoff, and (iv) if needed, adjust e-value cutoff and repeating the procedure. Since candidate sequence and HMM model quality varies, the best cutoff for each gene family differs from each other, and a rigorous manual validation is critical to ensure the quality of reference sequences. In addition, we clustered the reference sequences of each gene family into Functional Clusters (fClusters) based on the sequence similarity (95%), which allows further stratification of annotated reads by the same or highly related species.

**Figure 1.**
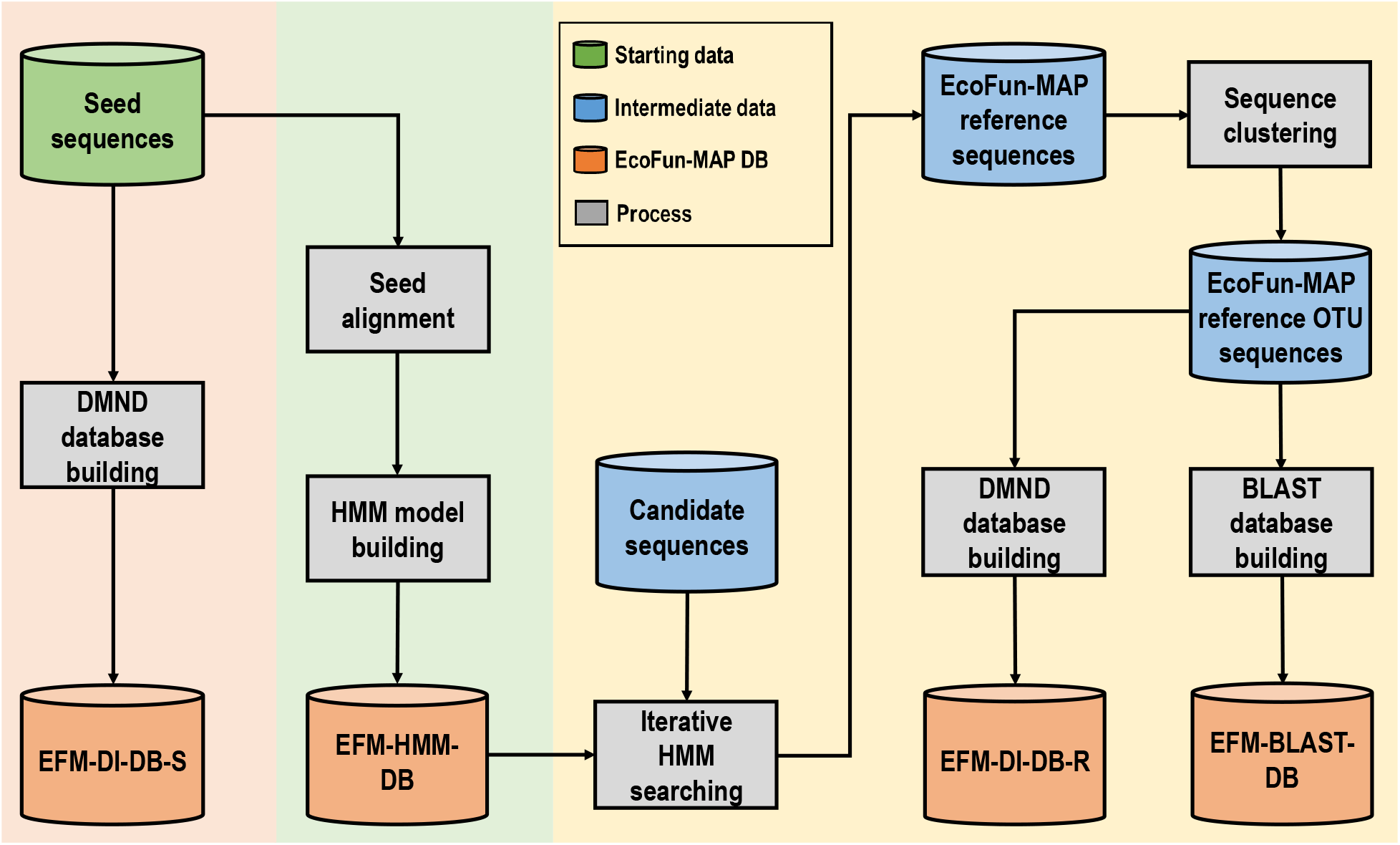
The flowchart of construction of databases/datasets in the development of EcoFun-MAP. Cylinders represent starting (green), intermediate (blue) and ending (orange) databases. Grey rectangles represent processing steps in construction, which take the content of databases or output of immediate upstream processing steps as input for processing. Four database modules have been constructed for EcoFun-MAP with the flowchart: a seed sequence-based DIAMOND index (EFM-DI-DB-S), Hidden Markov Models (HMMs) (EFM-HMM-DB), a functional gene reference sequence-based DIAMOND index (EFM-DI-DB-R) and a functional gene reference sequence based NCBI-BLAST index (EFM-BLAST-DB).

The reference database of EcoFun-MAP covered a total of 17 major functional gene categories and 160 primary subcategories (Table 1), which provides a comprehensive collection of functional genes that are important to biogeochemistry, ecology, environmental science, agriculture, and public health. For all 1,491 functional gene families included, we selected 14,500 seed sequences and built 1,862 HMM models. Meanwhile, a total of 1,217,363 reference functional gene sequences were retrieved and manually validated using the iterative HMM searching, which originated from about 50,000 taxonomical units that were distinguishable based on their taxonomical IDs. Based on these sequences, 280,247 fClusters were generated and further incorporated in EcoFun-MAP. Details about the coverage of EcoFun-MAP can be found in Table S1.

To fully take advantage of the reference database of EcoFun-MAP, we implemented a total of five workflows, which were labeled as ultra-fast, fast, moderate, conservative, and ultra-conservative (Fig. 2), respectively. All of the workflows used the same preprocessing procedure for quality trimming and gene prediction, in which the bases of low quality or ambiguity and excessively short reads were removed and gene fragments from qualified reads were identified. These workflows then diverge in the downstream analysis. In the ultra-fast workflow, preprocessed reads were directly searched against the EFM-DI-DB-S database. The fast workflow extended the ultra-fast workflow by further searching the EFM-DI-DB-S annotated reads against the EFM-HMM-DB and the EFM-BLAST-DB sequentially. Similarly, in the moderate workflow, the preprocessed reads were directly searched against the EFM-DI-DB-R. The conservative workflow extended the moderate by further searching the EFM-DI-DB-R annotated reads against the EFM-HMM-DB and then searching the resulting reads against the EFM-BLAST-DB. Finally, in the ultra-conservative workflow, the preprocessed reads were first searched against the EFM-HMM-DB, and then the resulting reads were searched against the EFM-BLAST-DB. In the end, all workflows provided an optional step to normalize counts of hits based on the average length of reference sequences from the gene families of the hits. Due to all these differences, the workflows should provide disparate performance in terms of both speed and accuracy, therefore allowing needed flexibility for data analysis in practice.

**Figure 2.**
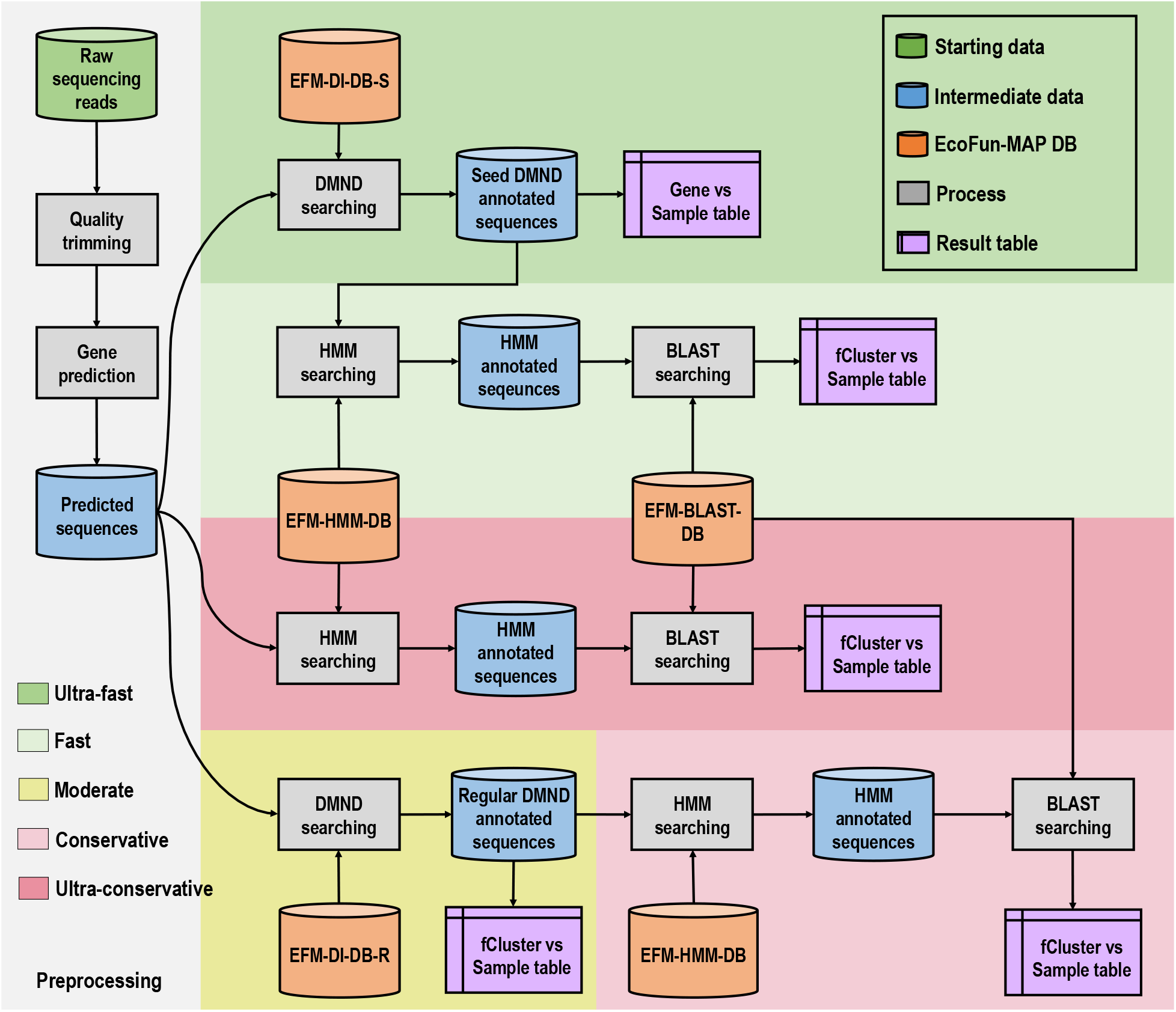
The flowchart of five workflows in EcoFun-MAP, which include ultra-fast (green background), fast (purple background), moderate (cyan background), conservative (yellowgreen background), and ultra-conservative (red background). The preprocessing steps are on the grey ground. Cylinders represent starting (green), intermediate (blue) and ending (orange) databases. Grey rectangles represent processing steps in construction, which take the content of databases or output of immediate upstream processing steps as input for processing. Shapes of yellow documents represent the resulting matrix-like table.

### Computing speed evaluation of EcoFun-MAP

To fully evaluate the computing speed of EcoFun-MAP in relation to input data size, we arbitrarily selected a groundwater sample (FW300), downsized it to subsamples of 0.7M, 3.5M, 7M, 35M and 70M reads, which accounted for ∼100M, ∼500M, ∼1G, ∼5G and ∼10G bases, and then ran all five EcoFun-MAP workflows on these subsamples. Each workflow was run with the same hardware configuration (10 nodes and 4 cores on each) and parameters. According to the design, we expected that the speed of the workflows should be ultra-fast > moderate > fast > conservative > ultra-conservative.

As expected, the ultra-fast workflow had the fastest speed, which was finished running on the largest subsample (70M reads) in 1,027 seconds (s), and then was followed by moderate (1,145 s), fast (1,506 s), conservative (1,865 s) and ultra-conservative (7,341 s) in order of decreasing speed (Table 2). The running of workflows on the largest subsample yielded the highest speed for all workflows (∼0.6-4.1M reads/min.), and the speed of workflows increased as the data size went up. The running on the smallest subsample yielded the lowest speed for all workflows (∼0.2-0.7M reads/min.). The ultra-fast workflow is >7 times faster compared to the ultra-conservative workflow for the largest subsample, but only 3 times faster for the smallest subsample in our test. Together these results suggest that EcoFun-MAP is fast (average speed from ∼0.4 to ∼2.5 M reads/min.) and highly scalable in high-throughput sequencing data analysis, in which time cost is expected to increase less than linearly as data size hikes, because of the increases of speed.

### Accuracy evaluation of EcoFun-MAP

Next, we evaluated the accuracy of EcoFun-MAP in terms of sensitivity and precision. Since the ground truth of gene annotation is not accessible for the metagenomes, direct estimation of accuracy is not possible. Here we used the annotation resulting from the ultra-conservative workflow as the ground truth to compare accuracy between the other four workflows, because the ultra-conservative workflow (i) performs homolog based search for every read, which takes into account information about protein domain structure and thus is considered to be more accurate than read mapping based only on sequence identity, and (ii) utilizes probabilistic models built on multiple sequence alignments and is thus generally more capable of detecting remote homologs than similarity search. By this definition, true positives (TP) of a workflow are the reads annotated by both the workflow and the ultra-conservative workflow; sensitivity is TP / total ultra-conservative annotations; precision is TP / total reads annotated by the evaluated workflow. We further defined precision and sensitivity at four category levels, based on TP reads within the same gene, secondary subcategory, primary subcategory or category as ultra-conservative annotations, respectively.

We ran all EcoFun-MAP workflows on the data from the 12 groundwater samples and compared their precision and sensitivity. The results (Table S2) showed that the numbers of hits produced by different workflows were ranged from ∼2.1 million (0.12%; moderate) to ∼81.1 million (4.46%; fast). Fast workflow produced the most hits of all (3.35-6.58%) across all samples, the moderate workflow produced the least (0.06-0.27%), and the conservative workflow had very similar yield (0.07-0.34%) as the ultra-conservative workflow (Table S2). For evaluated workflows, sensitivity rates (Table 3) were high in general (∼70% above). Fast workflow had the highest sensitivity rate at all levels (85.4-91.9%), which was then followed by conservative and ultra-fast workflow, and moderate workflow had the lowest (69.3%-69.8%) (Table 3). Differences of sensitivity rate across four annotation category levels (i.e., primary category, secondary category, gene family and gene) were small (< 0.5%) for moderate and conservative workflows, and higher in the ultra-fast (∼7.4%) and fast workflows (∼6.5%). Precision rate is the highest for moderate workflow at all levels (87.0-87.5%), but quite low for both fast (2.8-3.1%) and ultra-fast workflow (8.1-8.9%); with small variation across category levels (< 0.5%) for all workflows (Table 3). We note that the low precision of fast and ultra-fast workflow is mostly due to more reads were annotated by these two workflows, and those annotations not found by ultra-conservative workflow are not necessarily false positives. Together, the results suggest that EcoFun-MAP workflows should be chosen with consideration for distinct applications, e.g., fast and ultra-fast workflow for open gene search; conservative or ultra-conservative for stringent comparative analyses.

### Application to groundwater metagenomic analysis

To demonstrate the effectiveness of EcoFun-MAP in analyzing metagenomes, we ran all five workflows on a total of 12 groundwater samples collected from the Oak Ridge Integrated Field Research Challenge site^38,39^. These samples have labels including background (L0), low- (L1), intermediate- (L2), and high-contamination (L3), where L0 < L1 < L2 < L3 in terms of contamination level. SEED Subsystem annotation of these samples is also performed for comparison. Microbial community functional gene compositions were compared among the samples as shown in the DCA ordination plots (Figure. 4). The ordination results were consistent among all workflows. Samples from group L3 were observed to separate from other groups in all workflows with relatively high within-group distances. Clear separation of L2 samples from other groups was found in the moderate, conservative, and ultra-conservative workflows (Figure 4c, d and e). Clear separation of all four groups from each other was only observed in results based on the ultra-conservative workflow (Figure. 4e).

The functional gene richness from different workflows showed similar trends along the contamination gradient. The richness was significantly lower (p < 0.05) in L3 samples than in L0 samples, which was shown in analyses based on all EcoFun-MAP workflows and SEED Subsystem annotation. The analyses based on the fast workflow and SEED Subsystem^28^ annotation also showed a significantly lower (p < 0.05) richness of functional genes in L3 samples than in L2 samples. However, results from different workflows showed various estimations of richness changes. The ultra-fast and fast workflows estimated that richness of functional genes was ∼12% lower in L3 samples than in L1 samples, the moderate, conservative, and ultra-conservative workflows estimated that the richness of functional genes were ∼24% to ∼25% lower, and the SEED Subsystem annotation estimated that it was only ∼2.8% lower. Meanwhile, the fast workflow estimated that the richness of functional genes was ∼8.4% lower in L3 samples than in L2 samples, and SEED Subsystem annotation estimated ∼2.3% of lower richness. The results above suggest all workflows of EcoFun-MAP are capable of characterizing community-wide variations in groundwater metagenomes under the contamination gradient, with higher sensitivity compared to the SEED Subsystem annotations. This is probably because the SEED annotations include many universal physiological functions and genes which are less variable.

Next, we further analyzed relative abundances of major functional categories, including the category of C, N, S and P cycling, Metal homeostasis, Stress, Organic contaminant degradation, Antibiotic resistance, and Electron transfer, which are considered to be highly relevant to the study site, and compared them between different samples. The analysis was based on the ultra-conservative workflow. Relative abundances of functional genes from the C cycling category were lower in two of L3 samples (FW106 and FW021), which are two samples with the highest level of contamination in many heavy metals (e.g., Cr, Eu and Ce) (Figure S3), and those from the metal homeostasis category in the two samples were higher than other samples (Figure S4). Interestingly, sample FW104 from group L3, which had the highest level of Sulfate (SO_4_) of all samples (Figure S3), also has the highest relative abundance of S cycling genes (Figure S4). Response ratios (rr) of functional genes were calculated for comparing their relative abundances between sample group L0 and each of other groups (L1, L2 and L3). Among all genes with significant response ratios, we found abundances of homeostasis genes were significantly higher in L2 than L0 (*arrA* and *arxA;* rr=3.41 and 4.8) and in L3 than L0 (*corC, pcoA, mgtA* and *merP;* rr =1.09-5.38), and abundance of one C degradation gene (*ara*) was significantly lower (rr = -2.07) in L3 than L0 (Figure. 6). Meanwhile, a denitrification gene (*nirK*) was found to be more significantly abundant (rr = 1.76) in L3 than L0, which suggested a microbial response to higher nitrate concentrations in the L3 samples (Figure S3). Two oxygen-limitation-response genes, *narH* and *narJ*, from Stress category were more abundant (rr = 2.97 and 2.76) in L3 samples (DO=0.13-0.27) than L0 samples (DO=0.28-0.71), which suggested microbial response to low dissolved oxygen in highly contaminated wells.

## Discussion

EcoFun-MAP provides an efficient and accessible tool for analyzing shotgun metagenomic sequencing data from the perspective of ecological functions. With the typical speed of analysis from ∼0.6 to ∼4.1M reads/min, it helps overcome the computing barriers associated with deep functional profiling of microbes in a variety of environments.

The high computing efficiency of EcoFun-MAP is due to several reasons. First, the reference database is built, cleaned and optimized with a clear focus and much smaller in size compared to other general databases. With our curation efforts, the EcoFun-MAP database only has 1.5% of the size of NCBI RefSeq database (81,027,309 protein sequences; Mar 13th, 2017) while still provides a comprehensive coverage of keys genes from important ecological functions and geochemical processes. Such reduction strategy has been shown a useful solution for speeding up high-throughput sequencing data analysis^40^. Second, fast tools were selected for EcoFun-MAP and contributed substantially to the speed of EcoFun-MAP. For example, FragGeneScan+ used for gene prediction is 5-50 times faster than FragGeneScan at no cost of accuracy^41^. HMMER 3 is 100-1000 times faster than HMMER 2^42^. DIAMOND can be 20,000 times faster than BLASTX^23^. Third, EcoFun-MAP can process metagenomes in parallel and is deployed on an HPC cluster, which gains additional acceleration from advanced hardware. In addition to computing speed, EcoFun-MAP is also highly scalable, which is quite important since the volume of sequencing data continues to increase.

The reference database of EcoFun-MAP also has several unique features and advantages. First, the database has a clear microbial ecology focus compared to other recent tools annotating metagenomes with general metabolic genes, e.g., DRAM^43^ and METABOLIC^44^. The gene families were manually categorized into a hierarchical system that is similar as GeoChip^33–36^, which has been demonstrated consistently effective and easy to interpret in microbial ecology studies. Compared to FunGene^45^, a latest tool with a similar ecological focus, EcoFun-MAP covers 18 times more functional gene families as well as additional important function categories, e.g., stress response and virulence. Second, the reference sequences of each functional gene were clustered (fClusters), which provide a resolution beyond gene family and allow stratified analysis by groups of closely related microorganisms. In addition, EcoFun-MAP offers distinct database modules in a widely accepted format, e.g., HMM models. These modules enable speed and accuracy adjustment, underlie flexibility for different applications, and are easy to adapt for future tools and to extend for new genes and sequences. In the future, we will continue to maintain and update EcoFun-MAP databases as new knowledge (e.g. metagenome assembled genomes and genes^6,7,46–49^) comes in as well as exploring rapid algorithms (e.g. k-mer exact match^50–52^) for further speedup.

Apart from software tools like DRAM and METABOLIC, EcoFun-MAP is open for public use in the form of a website, so it is free of installation and configuration of dependencies or databases, and can be accessed using plain web browsers easily with Internet connection. While EcoFun-MAP was implemented and deployed based on advanced hardware and sophisticated bioinformatics tools, it requires little computer skills to use other than simple web-based user registration, uploading of datasets, and workflow selection or parameter setting. EcoFun-MAP is supported by an HPC infrastructure with fast CPUs, large memory, and hard disk space for public use. We consider this setup is ideal for data-intensive projects in microbial ecology and EcoFun-MAP should be highly accessible and usable to microbial ecologists in practice.

While the accuracy of EcoFun-MAP is difficult to directly evaluate, we adopted several ways to ensure that it is accurate. First, the reference database of EcoFun-MAP is rigorously curated, which ensures analysis quality at the beginning. Second, we used an iterative procedure to generate HMMs in EcoFun-MAP and manually tuned a key parameter (e.g., e-value cutoff of HMM search) per gene family, which should be more accurate compared to using an arbitrary or single universal parameter. In addition, EcoFun-MAP provides multiple predefined workflows accommodating disparate applications. The sensitivity is generally high for all workflows (∼70%). The ultra-fast and fast workflows showed low precision due to read identity-based searches, but are still useful for explorative analyses where detections with strong evidence are not mandatory. For example, we recommend the ultra-fast and fast workflow for discovering novel genes or gene fragments. Reassuringly, all EcoFun-MAP workflows revealed similar trends in the analysis of metagenomes from groundwater samples. Since the conservative workflow had both high sensitivity (∼85%) and precision (∼86%) rate, as well as speed (1.2M reads/min. on average), we set it to the default mode for EcoFun-MAP.

## Conclusion and availability

In this study, we developed EcoFun-MAP for functional analysis of shotgun metagenomic sequencing data from microbial ecology. EcoFun-MAP consists of references databases constructed with selective coverage of genes that are important to ecological functions, and multiple workflows for addressing disparate needs for speed and accuracy. Furthermore, EcoFun-MAP was implemented on the basis of High-Performance Computing (HPC) infrastructure with high accessible interfaces. Our analysis indicated that EcoFun-MAP is a fast and powerful pipeline for shotgun metagenome sequence data. EcoFun-MAP is open for public use and can be found available at our website: http://iegst1.rccc.ou.edu:8080/ecofunmap/.

## Material and Methods

### Selection of functional categories and genes

We limited the applicable scope of EcoFun-MAP to general microbial ecology studies and selected a total of 17 major categories (Table 1) of microbial genes that are associated with geochemical processes and ecological functions. These genes have been on functional gene arrays or GeoChip^33–36^, including Carbon (C), Nitrogen (N), Sulfur (S), and Phosphorus (P) cycling, antibiotic resistance, organic contaminant degradation, metal homeostasis, stress response, microbial defense, electron transferring, plant growth promotion, virulence, protists, viruses and others (metabolic pathways, pigment biosynthesis and *gyrB*). Then the functional genes are further divided into subcategories, yielding a three to four-level hierarchical organization: major category, primary subcategory, secondary category (optional), and functional gene. For example (Figure S1), the C cycling category (144 genes) consists of three primary subcategories, including C degradation (60 genes), C fixation (61 genes) and Methane (23 genes). The primary subcategory of C degradation has 18 secondary subcategories (e.g., Starch degradation, Cellulose degradation and Lignin degradation), the C fixation has 8 secondary subcategories (e.g., Calvin cycle, Dicarboxylate/4-hydroxybutyrate cycle and 3-hydroxypropionate bicycle), and the Methane has two secondary subcategories (i.e., Methane oxidation and Methanogenesis). Each secondary subcategory has the number of genes ranging from 1 to 21 (Figure S1).

### Retrieval of functional gene sequences

National Center for Biotechnology Information (NCBI) Entrez databases^53^ were used as the source to retrieve functional gene sequences for constructing EcoFun-MAP databases based on GeoChip databases. We manually crafted a keyword-based query for each functional gene, and submitted it programmatically to the Entrez databases to search and retrieve both protein and nucleotide candidate sequences via Entrez Programming Utilities (E-utilities)^53^. A typical search query is designed to consist of all aliases and variants names of the corresponding gene known to us, as well as other NCBI search constraints (e.g., organism), braces and logic operators (e.g., AND, OR and NOT). By carefully crafting the keyword-based query, the relevancy of research results can be improved as the number of the results drop, improving initial quality control before EcoFun-MAP database construction and reducing computational cost for later processing. For example, a keyword-based query for *nifH* gene (Suppl. Fig. 2) has returned 34,077 nucleotide records and 31,522 protein records, which were much less than 100,728 nucleotide records and 82,722 protein records in total returned by simply using “nifH” as the search query (retrieval test date: Jan. 23^rd^, 2017), and successfully excluded irrelevant records, such as *Sinorhizobium sp*. partial *nodA* gene (accession number: Z95242.1) and *Heliobacterium gestii* partial *anfH* gene (accession number: AB100834.1). Next, from records retrieved using keyword-based query search, a minimum of 1 to a few hundred seed sequences were selected manually on the basis of two criteria: (i) seed sequences must be experimentally confirmed in literature, and (ii) seed sequences must be distinctive from each other. Finally, redundant records (i.e., records with identical GenBank ID and description) were removed. To this end, candidate sequences and seed sequences have been prepared for each selected EcoFun-MAP gene and are ready for EcoFun-MAP database construction.

### Building EcoFun-MAP reference database

Reference database of EcoFun-MAP was built using the aforementioned candidate and seed sequences. The building process involves several key steps, including seed sequence alignment, HMM building, HMM searching, sequence clustering, DIAMOND index building and BLAST index building were implemented using ClustalW^54^, hmmbuild (HMMER3^42^), hmmsearch (HMMER3), CD-HIT^55^, DIAMOND and MAKEBLASTDB^37^, respectively. All of these tools were used with default parameters, except the CD-HIT used a customized threshold of clustering similarity at 95%.

### Design of EcoFun-MAP workflows

For the workflows, the key processing steps, including quality trimming, gene predicting, HMM searching, DIAMOND index searching and BLAST index searching were implemented using Btrim, FragGeneScan+, hmmsearch (HMMER3), DIAMOND and BLASTN, respectively. The workflows have preset parameters for each step, and can also accept users’ changes on the parameters for meeting specific speed or accuracy needs. For example, the Btrim used in the quality trimming for all the workflows has two major parameters: moving window size and average quality cutoff within the window. The default moving window size was set to 5 and the default average quality cutoff was set to 20 by EcoFun-MAP, but users can lower the moving window size or set higher the average quality cutoff to increase the quality of trimmed reads. All analyses in this study used preset parameters unless otherwise was mentioned.

### Deployment of EcoFun-MAP on HPC

The databases and workflows of EcoFun-MAP were deployed on an HPC cluster with a web-based Graphic User Interface (GUI) for access and job submission (Fig. 3). A single EcoFun-MAP job submission requires at the beginning a data file and all parameters that will be used for the selected workflow. EcoFun-MAP provides an FTP application for data file transferring and an HTTP application (website) to accept parameter settings. After being submitted, a job will be sent to the HPC cluster in a “first in, first out” (FIFO) order for further EcoFun-MAP processing. When executing a job, the HPC cluster will (i) break down the job into small pieces, (ii) map job pieces to available nodes, (iii) run the selected workflow for the pieces in parallel, and (iv) collect and reduce outputs of all pieces, and prepare final result for downloading by the job submitter. The implementation of EcoFun-MAP depends on both open-source software and in-house scripts. The FTP application was provided on the basis of installation and configuration of vsftpd (version 3.0.3). The parameter submission website was built using Django (version 1.11.5), In-house Perl, Python, Shell and SLURM job scheduling scripts were also used throughout EcoFun-MAP implementation. Their major functions or roles included the following: (i) job management, (ii) calling or executing bioinformatics tools, (iii) data file format conversion (e.g., convert FASTQ formatted file into a FASTA one), (iv) breaking down, mapping and reducing dataset and (v) data I/O and transferring. At last, the HPC cluster hosting EcoFun-MAP currently has two types (type I and type II) of computing nodes, and each type has 5 nodes, which consists of a total of 10 nodes for handling EcoFun-MAP tasks. The type I node has 24 cores and 64GB RAM, and the type II node has 24 cores and 128GB RAM. The HPC cluster also provides 128TB hard disk space for temporal storage of input, intermediate data and result from tasks of EcoFun-MAP.

**Figure 3.**
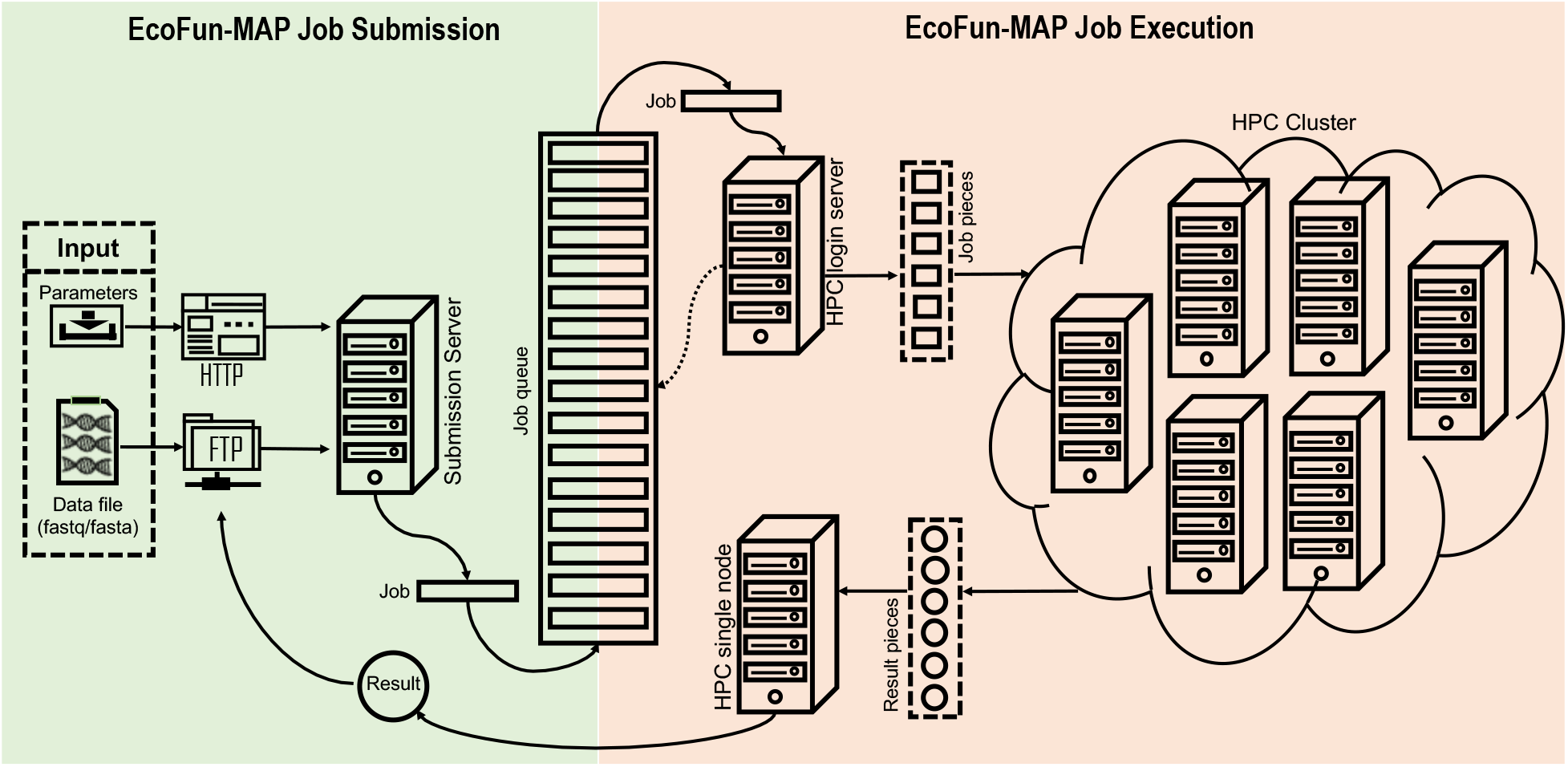
The scheme of implementation and deployment of EcoFun-MAP. Submissions of EcoFun-MAP jobs (green background) are handled by a standalone server. Further processing and execution of the jobs are performed on an HPC cluster.

**Figure 4.**
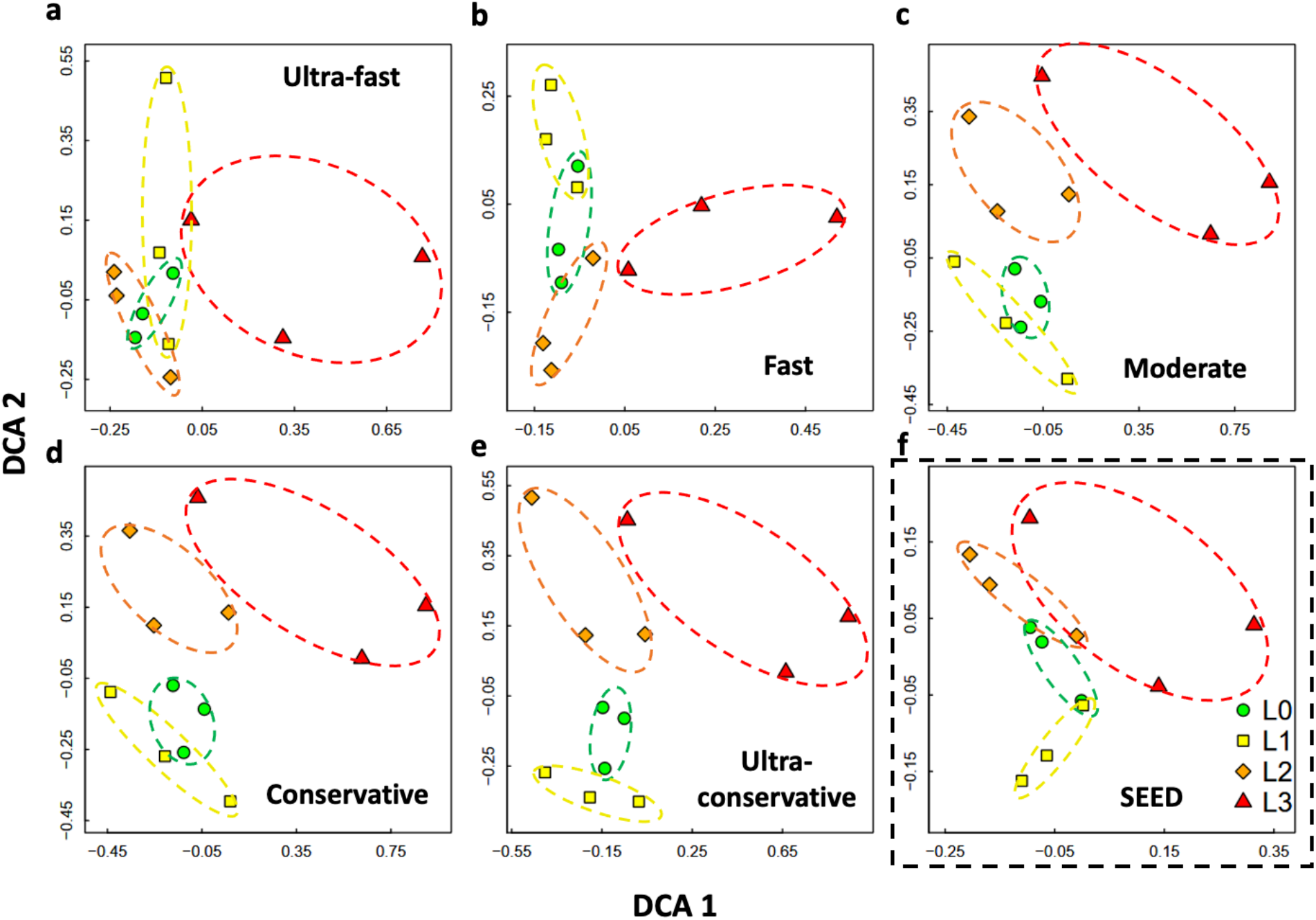
Detrended Correspondence Analysis (DCA) of functional gene compositions of metagenomes from 12 groundwater samples. Analyses of functional gene compositions based on results from five workflows of EcoFun-MAP are provided. Analysis based on the result from annotation based on SEED subsystem (boxed by dashed line) is also provided to contrast. Each sample is represented by a distinctive color. Cycles, squares, diamonds and triangles are used for showing samples from groups of L0, L1, L2 and L3, which are also cycled with green, yellow, orange and red eclipses, respectively.

**Figure 5.**
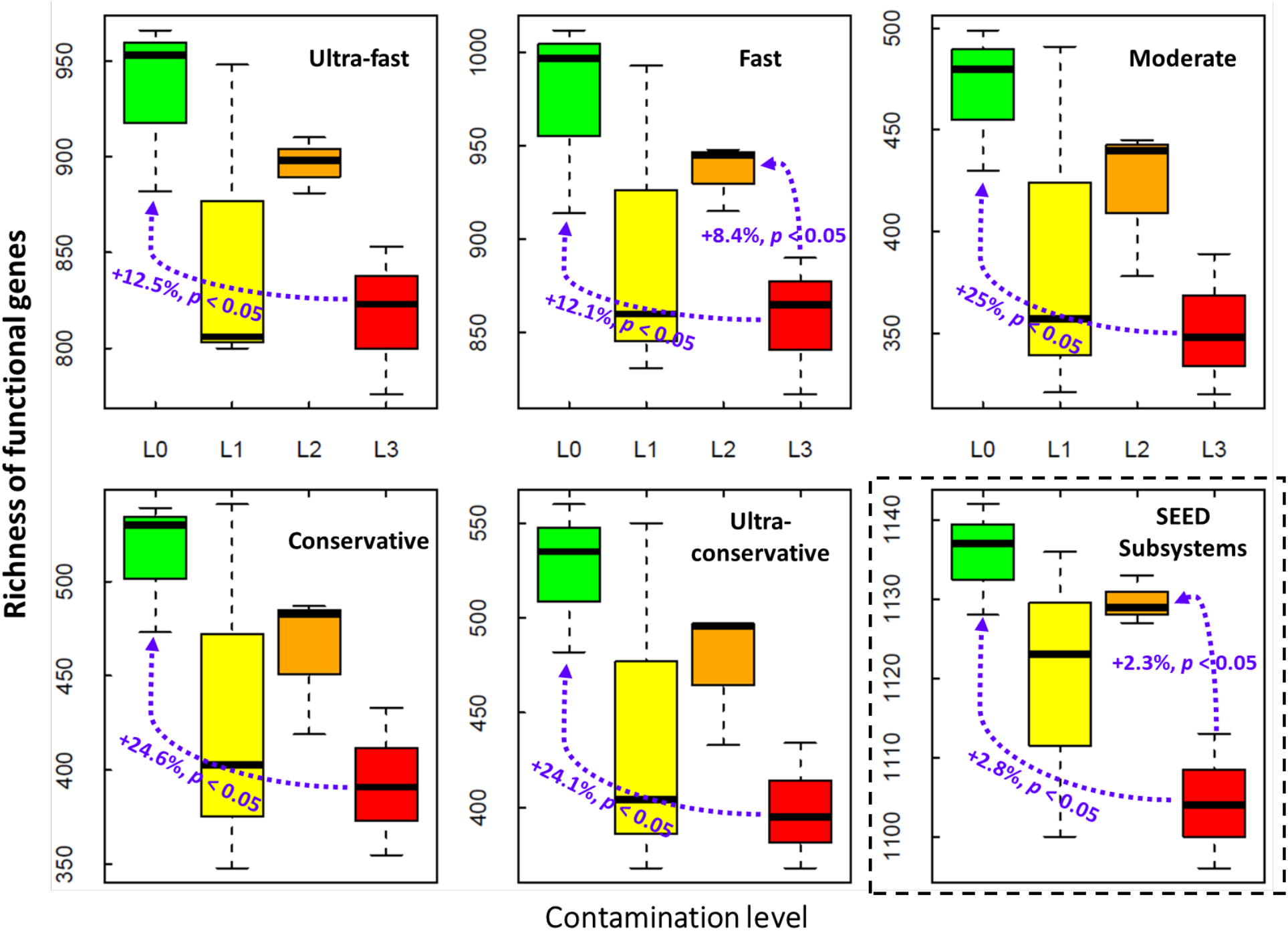
Richness of functional genes in metagenomes from 12 groundwater samples. A total of six boxplots show the richness of functional genes based on results from five workflows of EcoFun-MAP, as well as results from annotation based on SEED subsystem (boxed by dashed line). Boxes in color of green, yellow, orange and red are used for showing richness of functional genes for samples from groups of L0, L1, L2 and L3, respectively.

**Figure 6.**
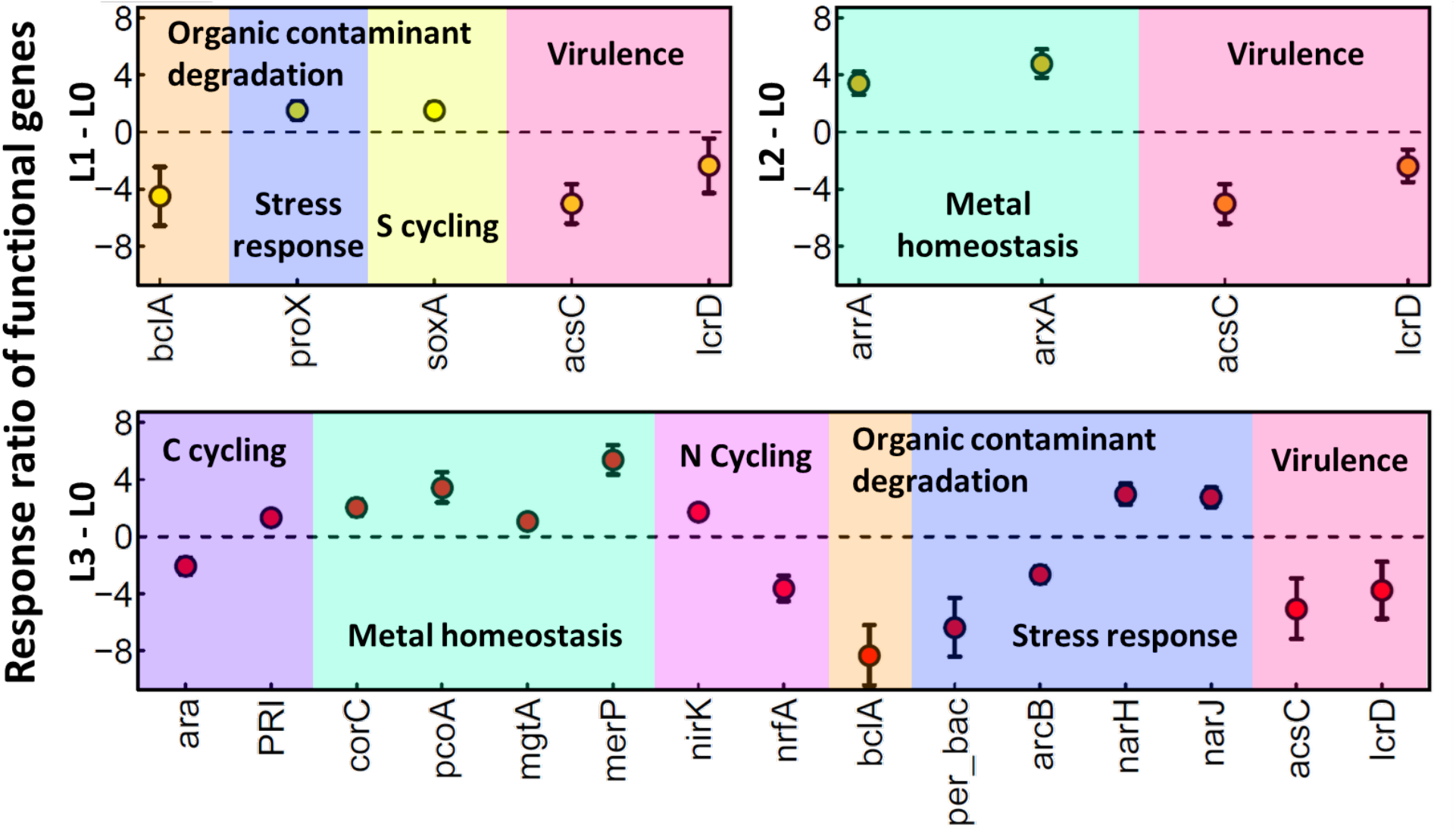
Response ratios of functional genes from comparisons between metagenomes from contaminated well samples and background well samples. Only significantly (*p value* < 0.05 in ANOVA followed by TukeyHSD) changed genes are included in the plot.

### Experimental datasets

Experimental datasets for showcasing and evaluating EcoFun-MAP were sequenced from groundwater samples from the Oak Ridge Integrated Field Research Challenge site^38^ (OR-IFRC; Oak Ridge, TN). The OR-IFRC site has gradients of salinity, pH and contaminants including Uranium, nitrate, sulfide, and other heavy metals^39,56^. In this study, 20 L groundwater was collected by 0.2-μm-pore-size filter from each of 12 locations under different contamination levels: background (L0), low- (L1), intermediate- (L2), and high-contamination (L3), with 3 samples for each level. Microbial community DNA was extracted from each sample using a modification of the Miller method^39,56,57^. The metagenome of each sample was sequenced using the shotgun method with HiSeq 3000 sequencer (Illumina, San Diego, CA). Upon completing HiSeq running, quality control was performed on the resulting raw reads. Duplicates and reads with ambiguous bases (>1) and poor-quality (average score <20) were discarded. Poor-quality bases (quality score <20) were trimmed. Finally, about 1,816.7 million of 150 bp reads were generated in total, which counted for about 272.5 Gbp data. The data size for each sample ranges from about 11.9 Gbp (GW199) to about 39.9 Gbp (FW300). More information about HiSeq output for each sample can be found in supplementary Table S2.

## Supporting information

main_and_suppl_tables

## Acknowledgements

The development of EcoFun-MAP was supported by the US Department of Energy, Office of Science, Genomic Science Program (Award Number: DE-SC0004601 and DE-SC0010715), and Office of Biological and Environmental Research’s (OBER) Biological Systems Research on the Role of Microbial Communities in Carbon Cycling program (Award number: DE-SC0004730 and DE-SC001057). The analysis of groundwater samples was supported by ENIGMA-Ecosystems and Networks Integrated with Genes and Molecular Assemblies (http://enigma.lbl.gov), a Scientific Focus Area Program at Lawrence Berkeley National Laboratory and is based upon work supported by the U.S. Department of Energy, Office of Science, Office of Biological & Environmental Research (contract number: DE-AC02-05CH11231). The development, implementation and maintenance of EcoFun-MAP by N.X. and D.N. were also partially supported by NSF Grants EF-2025558 and DEB-2129235.

## Figure Legends

**Figure S1.**
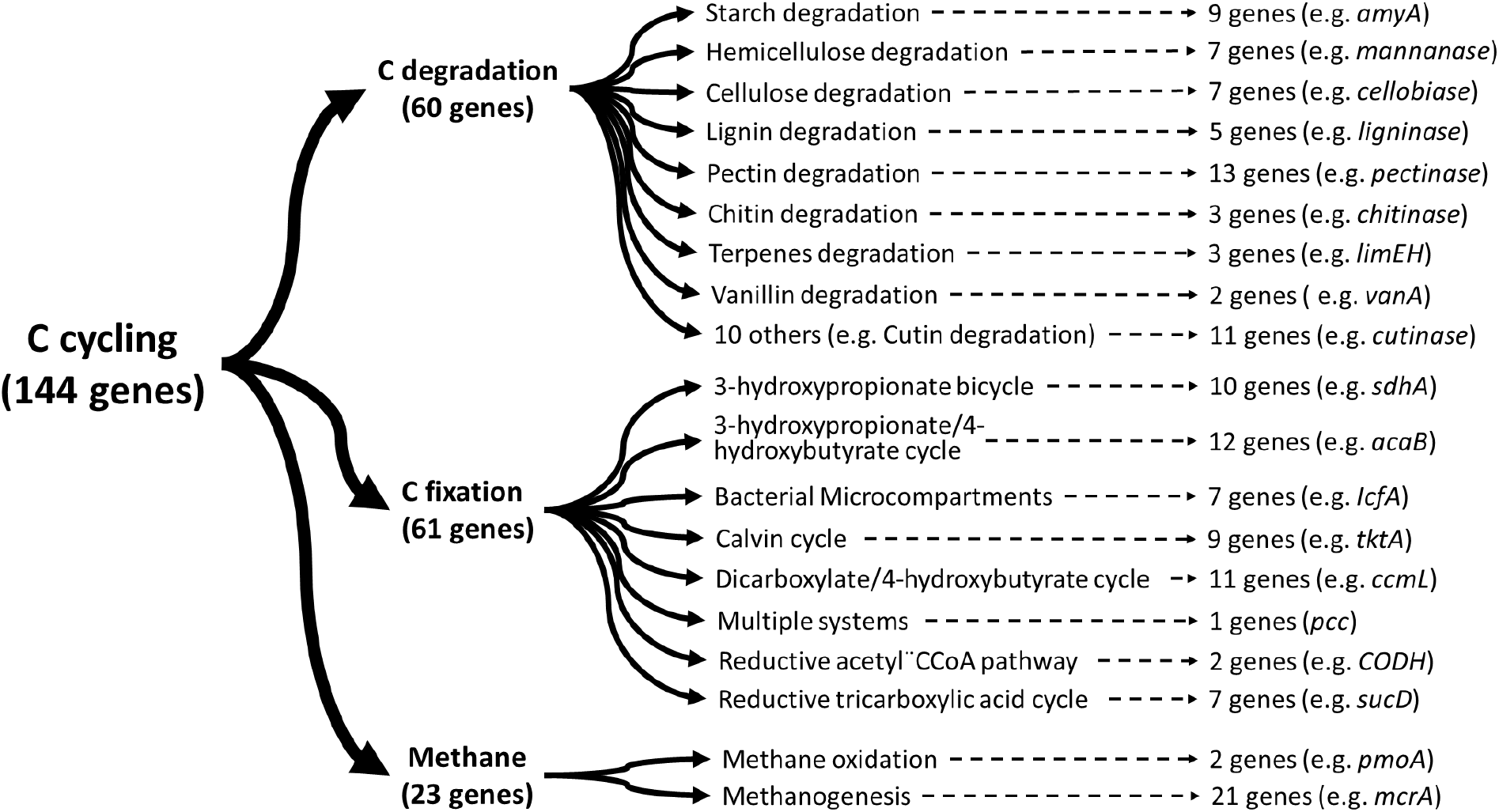
An example of organization of functional genes in EcoFun-MAP databases.

**Figure S2.**
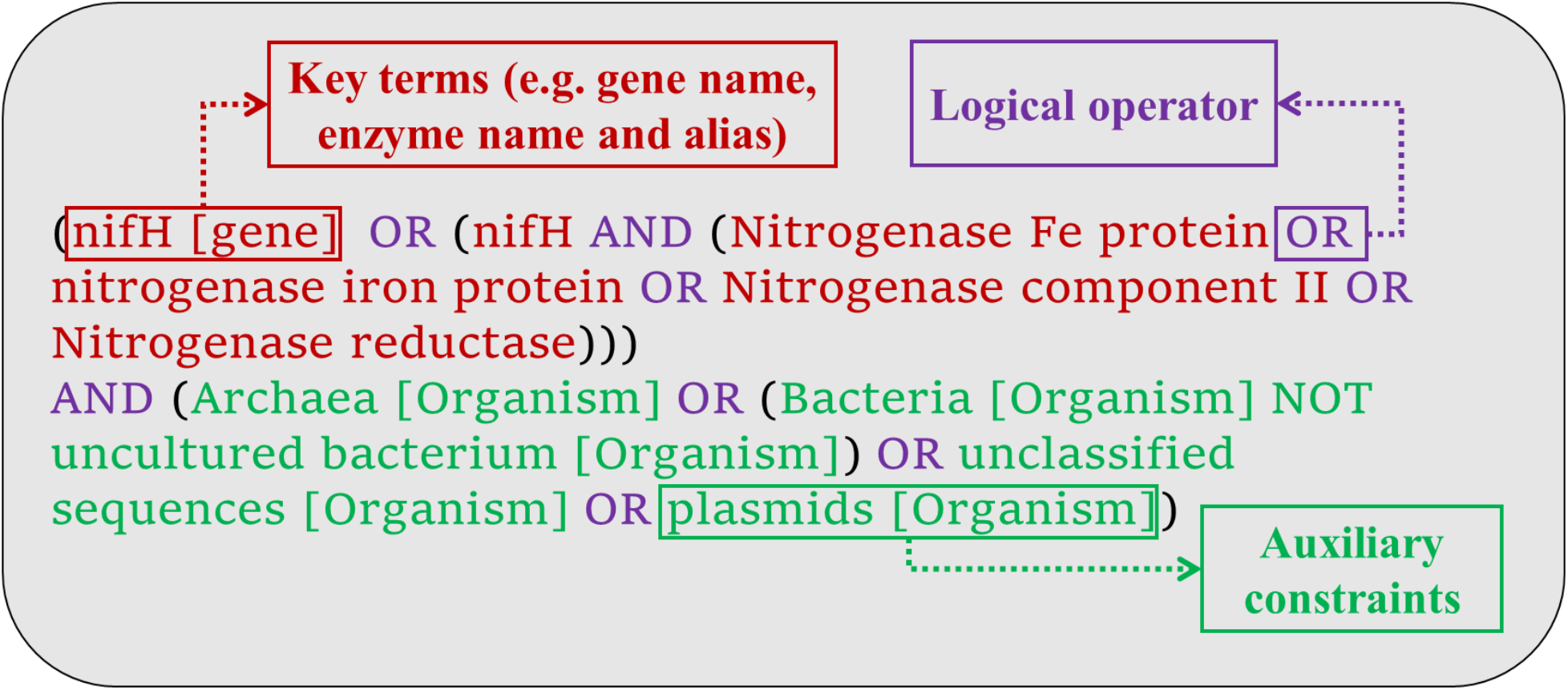
An example showing components that constitute a typical keyword query

**Figure S3.**
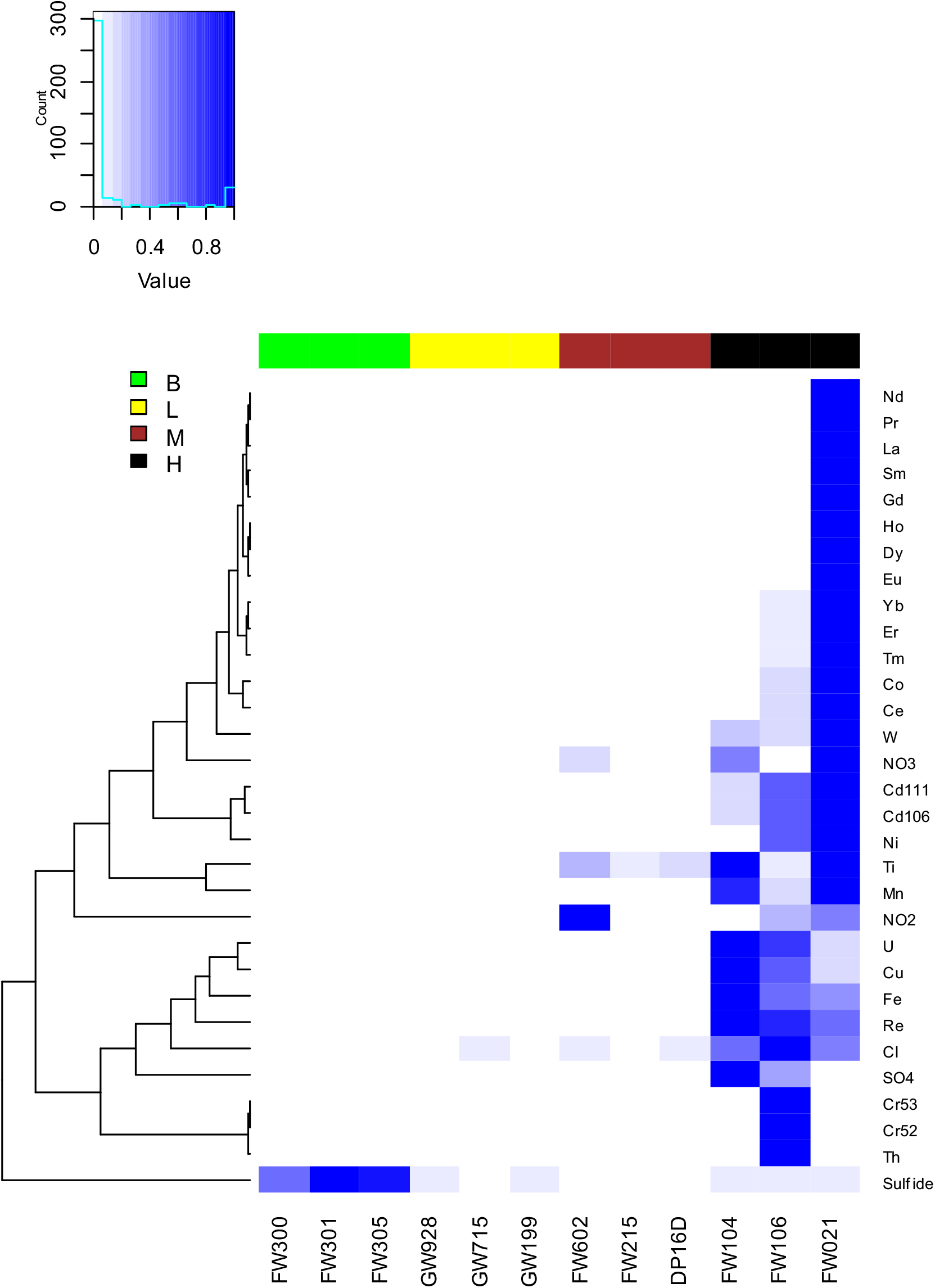
Heatmap showing the levels of measurements of environmental factors among 12 groundwater samples. The concentrations of pollutants were scaled to between 0 and 1 for a better visualization.

**Figure S4.**
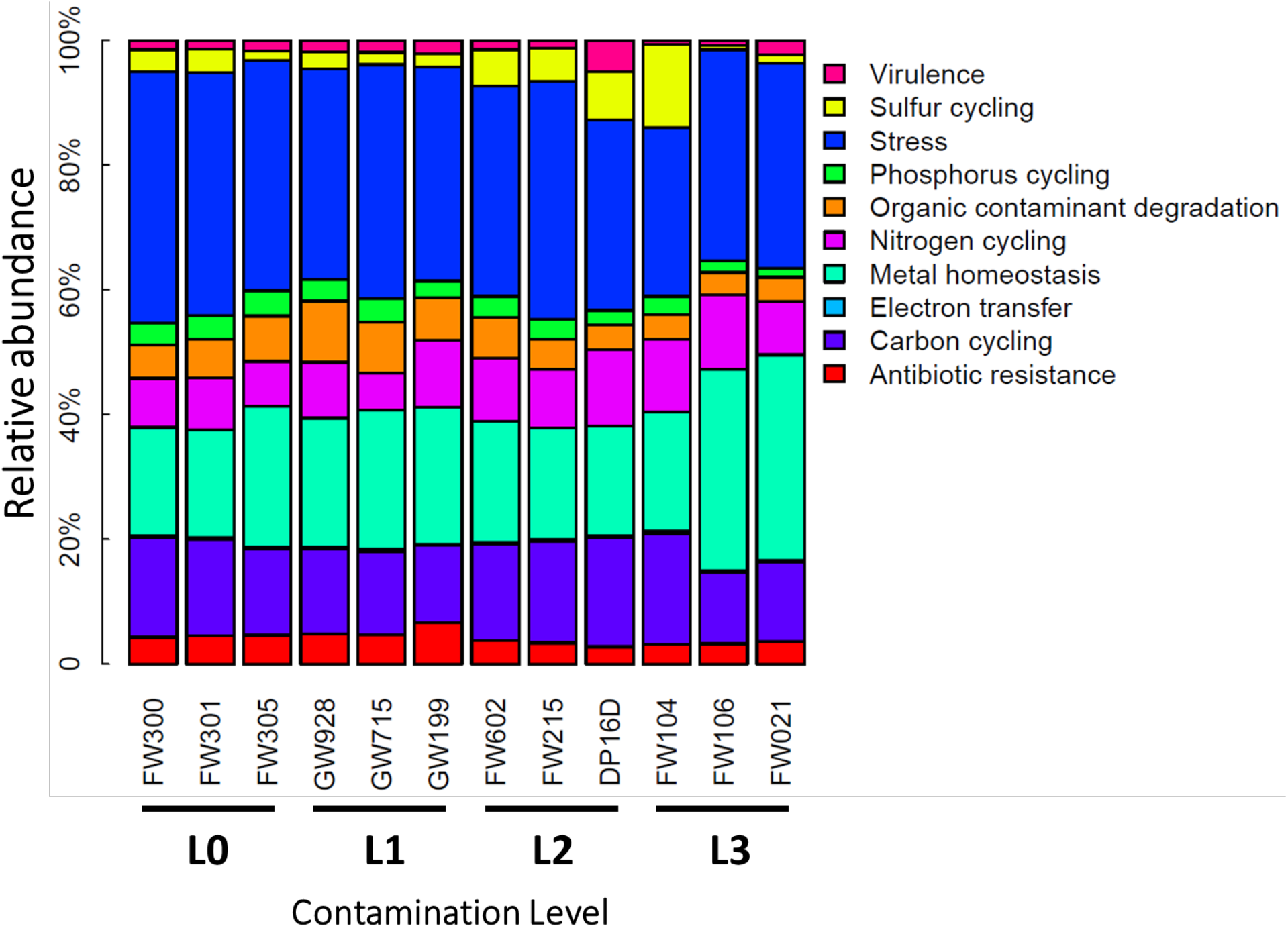
Relative abundances of selected major categories (based on result from Ultra-conservative workflow) in metagenomes from 12 groundwater samples.

