## Supplementary material for "EcoFun-MAP: An Ecological Function Oriented Metagenomic Analysis Pipeline": main_and_suppl_tables

### Table Legends

**Table 1.** Coverage of each functional category in the EcoFun-MAP database. * Only unqiue taxnomical IDs are counted**.**

**Table 2.** Time costs and speeds of five workflows in EcoFun-MAP. The evaluation was based on subsamples with different numbers of reads randomly drawn from the largest sample, the FW300. Preparing time refers to the time consumed outside the workflows, including file decompression, data transferring and partitioning and job scheduling.

**Table 3.** Sensitivity and precision of five workflows in EcoFun-MAP based on counts of hits from all samples.

**Table S1.** Coverage of functional genes in the EcoFun-MAP database.

**Table S2**. Metagenomic sequencing data size and hits of EcoFun-MAP analyses from the 12 groundwater samples.

**Table S3.** Sensitivity and precision of five workflows in EcoFun-MAP based on counts of hits from each sample.

**Table 1. Coverage of each functional category in the EcoFun-MAP database. * Only unqiue taxnomical IDs are counted.**

| Major categories | No. of primary subcategories | No. of genes | No. of seed sequences | No. of HMM models | No. of fClusters | No. of reference sequences | No. of covered taxonomical IDs* |
|---|---|---|---|---|---|---|---|
| C Cycling | 4 | 138 | 1,410 | 209 | 42,000 | 156,769 | 9,462 |
| N Cycling | 9 | 25 | 639 | 51 | 23,530 | 116,201 | 7,837 |
| S Cycling | 9 | 26 | 710 | 40 | 9,295 | 23,123 | 3,390 |
| P Cycling | 4 | 7 | 129 | 12 | 5,212 | 18,442 | 4,620 |
| Organic Contaminant Degradation | 9 | 149 | 1,290 | 250 | 19,139 | 81,479 | 7,755 |
| Antibiotic resistance | 3 | 19 | 234 | 48 | 24,450 | 105,376 | 6,708 |
| Stress | 24 | 100 | 2,529 | 159 | 47,809 | 223,627 | 9,379 |
| Metal Homeostasis | 25 | 120 | 658 | 161 | 5,529 | 247,826 | 7,217 |
| Microbial Defense | 2 | 65 | 495 | 66 | 10,613 | 33,499 | 3,204 |
| Metabolic Pathways | 2 | 4 | 54 | 8 | 2,110 | 8,338 | 2,210 |
| Plant Growth Promotion | 3 | 21 | 916 | 21 | 2,597 | 17,426 | 6,344 |
| Pigments | 6 | 26 | 128 | 29 | 1,755 | 4,398 | 1,024 |
| Electron transfer | 1 | 12 | 303 | 12 | 1,326 | 5,468 | 291 |
| Virulence | 42 | 608 | 3,467 | 613 | 10,415 | 63,605 | 4,463 |
| Virus | 3 | 113 | 984 | 113 | 8,079 | 63,324 | 15,697 |
| GyrB | 1 | 1 | 297 | 13 | 6,624 | 37,241 | 37,241 |
| Protist | 13 | 57 | 257 | 57 | 2,718 | 11,221 | 3,076 |
| **Total** | **160** | **1,491** | **14,500** | **1,862** | **280,247** | **1,217,363** | **49,018** |

**Table 2. Time costs and speeds of five workflows in EcoFun-MAP.** The evaluation was based on subsamples with different numbers of reads randomly drawn from the largest sample, the FW300. Preparing time refers to the time consumed outside the workflows, including file decompression, data transferring and partitioning and job scheduling.

| Mode | Preparing/main processing/total time (s) | | | | | Lowest/highest/average speed (No. of reads in M/min.) |
|---|---|---|---|---|---|---|
| | 0.7M reads (~100Mbp) | 3.5M reads (~500Mbp) | 7M reads (~1 Gbp) | 35M reads (~5 Gbp) | 70M reads (~10 Gbp) | |
| **ultra-fast** | 125/60/185 | 183/121/304 | 185/180/365 | 549/540/1,089 | 978/1,027/2,005 | ~0.7/4.1/2.5 |
| **fast** | 124/180/304 | 241/241/482 | 188/360/548 | 608/841/1,449 | 915/1,506/2,421 | ~0.2/2.8/1.5 |
| **moderate** | 159/60/219 | 186/180/366 | 190/240/430 | 615/602/1,217 | 862/1,145/2,007 | ~0.7/3.7/2.1 |
| **conservative** | 123/242/365 | 188/300/488 | 288/361/649 | 584/1,021/1,605 | 981/1,865/2,846 | ~0.2/2.2/1.2 |
| **ultra-conservative** | 125/181/306 | 187/485/672 | 189/840/1,029 | 617/3,966/4,583 | 920/7,341/8,261 | ~0.2/0.6/0.4 |

**Table 3. Sensitivity and precision of five workflows in EcoFun-MAP based on counts of hits from all samples.**

| Mode | Sensitivity rate | | | | Precision rate | | | |
|---|---|---|---|---|---|---|---|---|
| | Level 1 | Level 2 | Level 3 | Level 4 | Level 1 | Level 2 | Level 3 | Level 4 |
| **ultra-fast** | 70.2% | 73.8% | 74.5% | 77.6% | 8.1% | 8.5% | 8.6% | 8.9% |
| **fast** | **85.4%** | **88.3%** | **88.8%** | **91.9%** | 2.8% | 2.9% | 3.0% | 3.1% |
| **moderate** | 69.3% | 69.5% | 69.5% | 69.8% | **87.0%** | **87.2%** | **87.2%** | **87.5%** |
| **conservative** | 84.7% | 84.9% | 84.9% | 85.2% | 85.9% | 86.0% | 86.0% | 86.3% |

**Table S1. Coverage of functional genes in the EcoFun-MAP database. See spreadsheet.**

**Table S2. Metagenomic sequencing data size and hits of EcoFun-MAP analyses from the 12 groundwater samples.**

| Sample ID | Group label | Contamination level | No. of HiSeq reads (M)/ data amount (Gbp) | No. of hits (M)/percentage (%) | | | | |
|---|---|---|---|---|---|---|---|---|
| | | | | Ultra-fast | Fast | Moderate | Conservative | Ultra-conservative |
| FW300 | L0 | Background | ~266/39.9 | ~2.6/0.99 | ~8.9/3.35 | ~0.2/0.06 | ~0.2/0.07 | ~0.2/0.08 |
| FW301 | L0 | Background | ~164.5/25.7 | ~1.8/1.08 | ~5.9/3.62 | ~0.1/0.08 | ~0.2/0.09 | ~0.2/0.09 |
| FW305 | L0 | Background | ~97.2/14.6 | ~1.3/1.30 | ~4.5/4.63 | ~0.1/0.14 | ~0.2/0.18 | ~0.2/0.18 |
| GW199 | L1 | Low | ~79.6/11.9 | ~1.1/1.35 | ~3.5/4.41 | ~0.1/0.16 | ~0.2/0.19 | ~0.1/0.18 |
| GW928 | L1 | Low | ~92.5/13.9 | ~1.3/1.36 | ~4.3/4.61 | ~0.1/0.13 | ~0.1/0.16 | ~0.1/0.16 |
| GW715 | L1 | Low | ~202.3/30.3 | ~2.7/1.34 | ~9.6/4.76 | ~0.2/0.12 | ~0.3/0.16 | ~0.3/0.16 |
| DP16D | L2 | Medium | ~195.9/29.4 | ~2.4/1.23 | ~7.7/3.92 | ~0.1/0.07 | ~0.2/0.08 | ~0.2/0.08 |
| FW215 | L2 | Medium | ~171.3/25.7 | ~1.9/1.09 | ~6.0/3.52 | ~0.1/0.06 | ~0.1/0.07 | ~0.1/0.07 |
| FW602 | L2 | Medium | ~154.1/23.1 | ~2.0/1.29 | ~6.9/4.45 | ~0.2/0.12 | ~0.2/0.15 | ~0.2/0.15 |
| FW104 | L3 | High | ~94.5/14.2 | ~1.3/1.41 | ~4.8/5.05 | ~0.2/0.18 | ~0.2/0.21 | ~0.2/0.22 |
| FW106 | L3 | High | ~119.6/17.9 | ~2.1/1.74 | ~7.9/6.58 | ~0.3/0.27 | ~0.4/0.34 | ~0.4/0.35 |
| FW021 | L3 | High | ~178.9/26.8 | ~3.0/1.69 | ~11.1/6.20 | ~0.3/0.18 | ~0.4/0.22 | ~0.4/0.23 |
| **Total** | - | - | ~1816.7/272.5 | ~23.4/1.29 | ~81.1/4.46 | ~2.1/0.12 | ~2.7/0.15 | ~2.7/0.15 |

**Table S3. Sensitivity and precision of five workflows in EcoFun-MAP based on counts of hits from each sample.**

| Mode | Sample ID | Sensitivity rate | | | | Precision rate | | | |
|---|---|---|---|---|---|---|---|---|---|
| | | Level 1 | Level 2 | Level 3 | Level 4 | Level 1 | Level 2 | Level 3 | Level 4 |
| ultra-fast | FW300 | 71.5% | 74.0% | 74.9% | 78.2% | 5.5% | 5.7% | 5.8% | 6.0% |
| | FW301 | 72.5% | 74.7% | 75.6% | 78.8% | 6.2% | 6.4% | 6.4% | 6.7% |
| | FW305 | 68.8% | 70.8% | 71.7% | 74.7% | 9.5% | 9.8% | 9.9% | 10.4% |
| | GW199 | 72.9% | 74.7% | 75.2% | 77.8% | 9.6% | 9.8% | 9.9% | 10.2% |
| | GW928 | 70.1% | 72.1% | 72.9% | 76.0% | 8.2% | 8.5% | 8.6% | 8.9% |
| | GW715 | 67.1% | 69.3% | 70.2% | 73.9% | 8.2% | 8.4% | 8.5% | 9.0% |
| | DP16D | 70.9% | 73.3% | 73.8% | 76.3% | 4.8% | 5.0% | 5.0% | 5.2% |
| | FW215 | 73.5% | 76.1% | 76.7% | 79.7% | 4.9% | 5.0% | 5.1% | 5.3% |
| | FW602 | 70.9% | 73.5% | 74.3% | 77.8% | 8.4% | 8.7% | 8.8% | 9.2% |
| | FW104 | 73.4% | 77.7% | 78.0% | 80.6% | 11.3% | 12.0% | 12.0% | 12.4% |
| | FW106 | 68.3% | 74.4% | 74.7% | 77.6% | 13.8% | 15.0% | 15.0% | 15.6% |
| | FW021 | 69.6% | 75.5% | 76.3% | 79.9% | 9.4% | 10.1% | 10.3% | 10.7% |
| fast | FW300 | 85.1% | 87.4% | 87.8% | 91.2% | 1.9% | 2.0% | 2.0% | 2.1% |
| | FW301 | 86.4% | 88.4% | 88.9% | 92.0% | 2.2% | 2.2% | 2.3% | 2.3% |
| | FW305 | 86.8% | 88.8% | 89.3% | 92.4% | 3.4% | 3.5% | 3.5% | 3.6% |
| | GW199 | 87.3% | 88.6% | 88.9% | 91.1% | 3.5% | 3.6% | 3.6% | 3.7% |
| | GW928 | 86.6% | 88.3% | 88.7% | 91.8% | 3.0% | 3.1% | 3.1% | 3.2% |
| | GW715 | 85.8% | 87.9% | 88.4% | 92.1% | 2.9% | 3.0% | 3.0% | 3.2% |
| | DP16D | 85.0% | 86.7% | 86.8% | 89.4% | 1.8% | 1.8% | 1.8% | 1.9% |
| | FW215 | 86.0% | 88.1% | 88.4% | 91.6% | 1.8% | 1.8% | 1.8% | 1.9% |
| | FW602 | 84.9% | 86.9% | 87.4% | 91.0% | 2.9% | 3.0% | 3.0% | 3.1% |
| | FW104 | 86.7% | 88.4% | 88.8% | 91.6% | 3.7% | 3.8% | 3.8% | 3.9% |
| | FW106 | 83.9% | 88.9% | 89.6% | 92.8% | 4.5% | 4.7% | 4.8% | 4.9% |
| | FW021 | 84.3% | 89.4% | 90.2% | 93.2% | 3.1% | 3.3% | 3.3% | 3.4% |
| moderate | FW300 | 69.0% | 69.2% | 69.2% | 69.5% | 86.1% | 86.3% | 86.3% | 86.7% |
| | FW301 | 70.1% | 70.2% | 70.2% | 70.5% | 84.1% | 84.3% | 84.3% | 84.6% |
| | FW305 | 66.5% | 66.7% | 66.7% | 67.0% | 86.2% | 86.5% | 86.5% | 86.9% |
| | GW199 | 71.1% | 71.2% | 71.2% | 71.5% | 78.3% | 78.5% | 78.5% | 78.8% |
| | GW928 | 68.0% | 68.1% | 68.1% | 68.5% | 84.7% | 84.8% | 84.8% | 85.3% |
| | GW715 | 65.3% | 65.4% | 65.5% | 65.8% | 88.0% | 88.2% | 88.2% | 88.7% |
| | DP16D | 69.0% | 69.1% | 69.1% | 69.3% | 82.4% | 82.6% | 82.6% | 82.8% |
| | FW215 | 71.5% | 71.7% | 71.7% | 72.0% | 82.9% | 83.1% | 83.1% | 83.4% |
| | FW602 | 69.6% | 69.7% | 69.8% | 70.1% | 88.7% | 88.9% | 88.9% | 89.3% |
| | FW104 | 73.2% | 73.3% | 73.3% | 73.5% | 87.1% | 87.2% | 87.2% | 87.4% |
| | FW106 | 69.4% | 69.6% | 69.6% | 69.8% | 91.1% | 91.4% | 91.4% | 91.7% |
| | FW021 | 70.8% | 71.0% | 71.0% | 71.3% | 90.5% | 90.7% | 90.8% | 91.1% |
| conservative | FW300 | 82.9% | 83.1% | 83.1% | 83.4% | 85.2% | 85.3% | 85.3% | 85.7% |
| | FW301 | 84.4% | 84.5% | 84.5% | 84.8% | 84.1% | 84.2% | 84.2% | 84.5% |
| | FW305 | 84.2% | 84.4% | 84.4% | 84.7% | 84.7% | 84.9% | 85.0% | 85.3% |
| | GW199 | 84.2% | 84.3% | 84.3% | 84.5% | 79.0% | 79.1% | 79.1% | 79.4% |
| | GW928 | 83.8% | 83.9% | 83.9% | 84.3% | 84.0% | 84.1% | 84.1% | 84.4% |
| | GW715 | 83.6% | 83.8% | 83.8% | 84.2% | 85.6% | 85.8% | 85.8% | 86.2% |
| | DP16D | 82.5% | 82.6% | 82.6% | 82.8% | 83.1% | 83.2% | 83.2% | 83.5% |
| | FW215 | 84.0% | 84.2% | 84.2% | 84.5% | 83.3% | 83.5% | 83.5% | 83.7% |
| | FW602 | 84.4% | 84.5% | 84.5% | 84.9% | 87.2% | 87.4% | 87.4% | 87.7% |
| | FW104 | 85.6% | 85.7% | 85.7% | 85.9% | 86.7% | 86.8% | 86.8% | 87.0% |
| | FW106 | 86.8% | 86.9% | 87.0% | 87.2% | 88.7% | 88.9% | 88.9% | 89.1% |
| | FW021 | 86.3% | 86.4% | 86.4% | 86.7% | 88.7% | 88.9% | 88.9% | 89.2% |
